# Deep sequencing of yeast and mouse tRNAs and tRNA fragments using OTTR

**DOI:** 10.1101/2022.02.04.479139

**Authors:** H. Tobias Gustafsson, Carolina Galan, Tianxiong Yu, Heather E. Upton, Lucas Ferguson, Ebru Kaymak, Zhiping Weng, Kathleen Collins, Oliver J. Rando

## Abstract

Among the major classes of RNAs in the cell, tRNAs remain the most difficult to characterize via deep sequencing approaches, as tRNA secondary structure and nucleotide modifications can both interfere with cDNA synthesis by commonly-used reverse transcriptases (RTs). Here, we benchmark a recently-developed RNA cloning protocol, termed Ordered Two-Template Relay (OTTR), to characterize intact tRNAs and tRNA fragments in budding yeast and in mouse tissues. We find that OTTR robustly captures full-length tRNAs in budding yeast and in mouse testis, with relatively low levels of premature termination at known barriers – 1-methylguanine, N2,N2-dimethylguanine, and 1-methyladenine – to typical reverse transcriptases. Moreover, these and several other nucleotide modifications leave misincorporation signatures in OTTR datasets which enables their detection without any additional protocol steps. Turning to analysis of small RNAs such as tRNA cleavage products, we compare OTTR with several standard small RNA-Seq protocols, finding that OTTR provides the most accurate picture of tRNA fragment levels by comparison to “ground truth” Northern blots. Applying this protocol to mature mouse spermatozoa, our data dramatically alter our understanding of the small RNA cargo of mature mammalian sperm, revealing a far more complex population of tRFs – including both 5’ and 3’ tRNA halves derived from the majority of tRNAs – than previously appreciated. Taken together, our data confirm the superior performance of OTTR to commercial protocols in analysis of tRNA fragments, and force a reappraisal of potential epigenetic functions of the sperm small RNA payload.

## INTRODUCTION

tRNAs represent the physical embodiment of the genetic code, and are broadly expressed in all cell types in the body and across a wide range of environmental conditions. Nonetheless, there is increasing evidence that the cellular repertoire of tRNAs differs between different cell types, and within a given cell type can be shaped by external factors from proliferation rate [1, 2] to metabolite levels [3]. Mature tRNAs are also cleaved in response to cellular stressors [4-7], and the resulting cleavage products – broadly known as tRNA fragments, or tRFs – are increasingly appreciated as potential regulatory molecules in their own right [8-10].

In contrast to most other RNA species, characterization of tRNA and tRF levels by deep sequencing has been hampered by technical difficulties in synthesis of cDNA. tRNAs are subject to a wide range of covalent nucleotide modifications, with some ∼15-20% of all tRNA nucleotides thought to be covalently modified to form species ranging from 5-methylcytosine and pseudouridine to more complex modifications like wybutosine or methoxy-carbonyl-methyl-thiouridine [11-13]. Several of these modifications, particularly N1-methylguanosine (m^1^G), N1-methyladenosine (m^1^A), N3-methylcytosine (m^3^C), and N2,N2-dimethylguanosine (m^2^_2_G), are known to interfere with commonly-used reverse transcriptases and prevent the synthesis of full-length cDNAs. As a result, until recently systematic analyses of intact tRNA levels have typically relied on microarray hybridization [14, 15] to avoid a requirement for reverse transcription. With respect to tRNA fragments, although typical deep sequencing efforts do capture some tRNA cleavage products, they are clearly limited to only a subset of the tRFs in a given sample. For instance, a large number of groups have characterized small (18-40 nt) RNAs in mammalian sperm, with most such studies documenting very high levels of 5’ fragments of a small handful of tRNAs (most notably including Gly-GCC, Glu-CTC, and Val-CAC). Yet Northern blots show that 3’ tRNA fragments are also present in these samples [16, 17], but are not captured by typical commercial library preparation protocols such as Illumina TruSeq.

A number of methods have been developed in the past few years to enable analysis of intact tRNAs by deep sequencing. For instance, to reduce barriers to reverse transcription caused by secondary structures, Hydro-tRNAseq introduced limited hydrolysis of full-length tRNAs to yield short fragments for cloning and sequencing [18-20]. Alternatively, several early protocols leveraged the highly processive thermostable group II intron reverse transcriptase (TGIRT) to overcome tRNA secondary structures [21], along with enzymatic demethylation of m^1^G, m^1^A, and m^3^C by bacterial AlkB in an attempt to avoid premature RT termination [22-24]. Although these first-generation protocols yielded few full length tRNA sequences, a substantially-improved TGIRT-based protocol – mim-tRNAseq – was recently shown to efficiently capture full length tRNAs with only modest levels of premature RT termination [25]. Other recently-developed tRNA cloning protocols include YAMAT-Seq [26] and QuantM-Seq [27].

Most recently, Collins and colleagues developed a novel protocol – Ordered Two-Template Relay, or OTTR [28] – based on an engineered version of the *B. Mori* R2 retroelement reverse transcriptase [29]. Benchmarking a large number of protocols against a defined mixture of small RNAs revealed significantly lower cloning bias for OTTR than for any commercial protocol [28]. Moreover, characterization of RNA in tissue culture cell lines by OTTR revealed substantial levels of intact tRNAs, suggesting that OTTR could be a promising protocol for tRNA sequencing applications.

Here, we set out to explore the utility of OTTR for analysis of intact tRNAs and tRNA fragments in several biological systems. We successfully sequenced full length intact tRNAs from budding yeast and from mouse testis, and confirmed that a number of specific nucleotide modifications induce mismatch signatures in the tRNA sequencing dataset. We next turned to analysis of small RNA populations in three systems: budding yeast overexpressing the RNaseT2 family member RNY1p, mouse cauda epididymis, and mature cauda epididymal sperm. Comparison of OTTR with several commercial protocols, coupled with gold standard Northern blot validation, confirmed that OTTR more accurately captures tRNA fragments than either NEB Next, Illumina Truseq, or a typical in-house protocol based on adaptor ligation. In the mouse samples, we show that OTTR captures a far greater variety of tRNA cleavage products, including abundant 3’ tRNA fragments, that are invisible to all other protocols examined. Taken together, our data provide an updated view of the mouse sperm small RNA payload, and highlight the utility of OTTR for analysis of tRNAs and tRNA fragments by deep sequencing.

## RESULTS

### Cloning of full-length tRNAs in budding yeast and mouse testis

We initially sought to compare OTTR with several commercial protocols for analysis of small (18-40 nt) RNA populations in mouse sperm. However, our proof of concept datasets revealed abundant nucleotide mismatches at presumed sites of tRNA nucleotide modifications (see below), as observed in multiple prior deep sequencing analyses of tRNAs [22-25]. We therefore set out first to sequence intact tRNAs to empirically characterize effects of nucleotide modifications on deep sequencing libraries, to help guide bioinformatic analyses of tRNA-derived sequences.

We focused on two biological systems. First, given the ease of growing large quantities of *S. cerevisiae* for validation of our sequencing data by Northern blots, along with the extensive characterization of tRNA modifications in this species, we sequenced intact tRNAs from actively growing budding yeast. Second, mouse sperm are thought to carry a payload of small RNAs dominated by 5’ tRNA halves [30], and an increasing number of studies have implicated mouse sperm RNAs as potential mediators of intergenerational paternal effects. However, given that sperm do not carry substantial levels of intact tRNAs [16, 17], we instead first turned to mouse testis samples as a source of intact mammalian tRNAs.

For each sample, we generated total RNA, then either fractionated RNAs over a spin column to enrich for <200 nt RNAs (**Supplemental Figure S1**), or gel-purified 60-100 nt RNAs. A second size selection step was added following cDNA synthesis, to deplete adaptor dimers and enrich for libraries carrying ∼60-100 bp inserts. Resulting OTTR libraries were then sequenced to an average of ∼10 million reads. Surprisingly, we consistently recovered more full-length tRNAs when using the more lenient RNA sizing by *mir*Vana spin column (**Figure 1A**), suggesting that the process of gel-mediated size selection likely results in tRNA fragmentation. We therefore focused downstream analyses on tRNA-mapping reads in the *mir*Vana-sized OTTR libraries.

**Figure 1.**
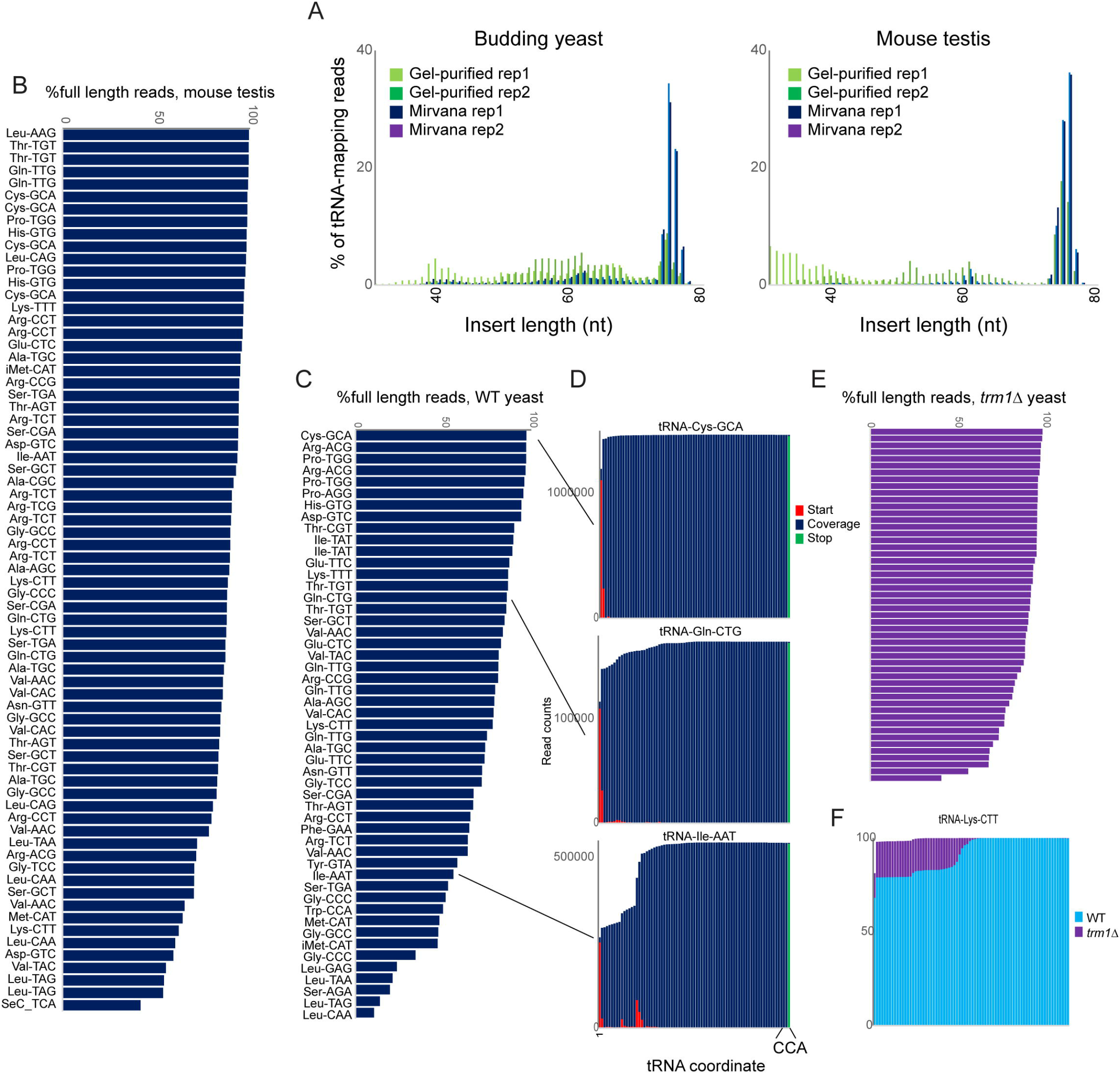
OTTR successfully captures full length tRNAs in yeast and mouse. A) Insert length distributions for full-length tRNA OTTR libraries for budding yeast, and mouse testis, as indicated. Libraries were prepared following one of two initial size selection steps: “Gel” refers to libraries build using total RNA subject to acrylamide gel-based purification of 60-100 nt RNAs, “Mirvana” refers to libraries build using the small (<200 nt) fraction (**Supplemental Figure S1**) recovered from *mir*Vana RNA spin columns. B-C) Efficient capture of full length tRNAs using OTTR. For each tRNA species with over 1000 reads, percentage of full length tRNA reads was calculated. For each species, mouse (B) and yeast (C), tRNAs are ordered from highest to lowest coverage. D) Coverage plots for three exemplar tRNAs in the yeast OTTR dataset. Red and green bars show sequence starts and stops, respectively, while blue bars show sequence coverage. E-F) Improved full-length tRNA coverage in *trm1*Δ yeast lacking m^2^_2_G. As in panels (C-D).

Initial mapping of OTTR reads to mature tRNA sequences using standard analytical pipelines was hindered by the high numbers of sequence mismatches likely driven by modified nucleotides in tRNAs. We therefore turned to the tRAX analytical pipeline [31], a mismatch-tolerant pipeline that accounts for the wide range of post-transcriptional modifications to tRNAs that can complicate typical RNA mapping pipelines. For each tRNA we calculated the percentage of full length reads, finding that the majority of tRNAs exhibited only modest levels (∼10-30% of reads) of premature RT termination in mouse and yeast (**Figures 1B-D**). Comparison to a range of prior tRNA sequencing datasets [20, 22, 23, 25-27] confirmed that OTTR was comparable to mim-tRNAseq and YAMAT-Seq in terms of capturing full-length tRNAs, while the remaining protocols all exhibited much more extensive premature RT termination (**Supplemental Figure S2**).

Visualization of sequence coverage and read start and stop locations for several tRNAs (**Figure 1D**) revealed that a subset of premature RT termination events occurred at known nucleotide modification sites including the common m^1^G modification found at position 9 of many tRNAs (eg, tRNA-Ile-AAT). To directly test whether premature RT termination is affected by nucleotide modifications, we prepared full-length tRNA libraries from *trm1*Δ yeast lacking the methylase responsible for m^2^_2_G [32]. We find further gains in the efficiency of full-length tRNA capture in this strain background (**Figures 1E-F**), suggesting that this nucleotide modification presents a partial barrier to reverse transcription.

### Signatures of nucleotide modifications in full-length tRNA sequences

Many of the nucleotide modifications in tRNAs involve chemical alterations that affect the pattern of hydrogen bond donors and acceptors at the base pairing interface. As a result, “incorrect” nucleotides (relative to those expected from the tRNA’s genomic sequence) can be incorporated into cDNA at these positions during the process of reverse transcription, resulting in a “mutation/misincorporation” signature for modified nucleotides in deep sequencing data.

Examination of individual tRNAs revealed high levels of misincorporation in yeast at multiple positions throughout the tRNA (**Figure 2A**), with mismatches localized at various known modification sites including the expected mismatches at known m^1^G, m^2^_2_G, m^3^C, and m^1^A nucleotides. Examination of the same tRNA species in our *trm1*Δ dataset revealed the expected loss of nucleotide misincorporation at G26 in this mutant (**Figure 2B**), confirming that the m^2^_2_G nucleotide modification is responsible for the mutational signature at this position.

**Figure 2.**
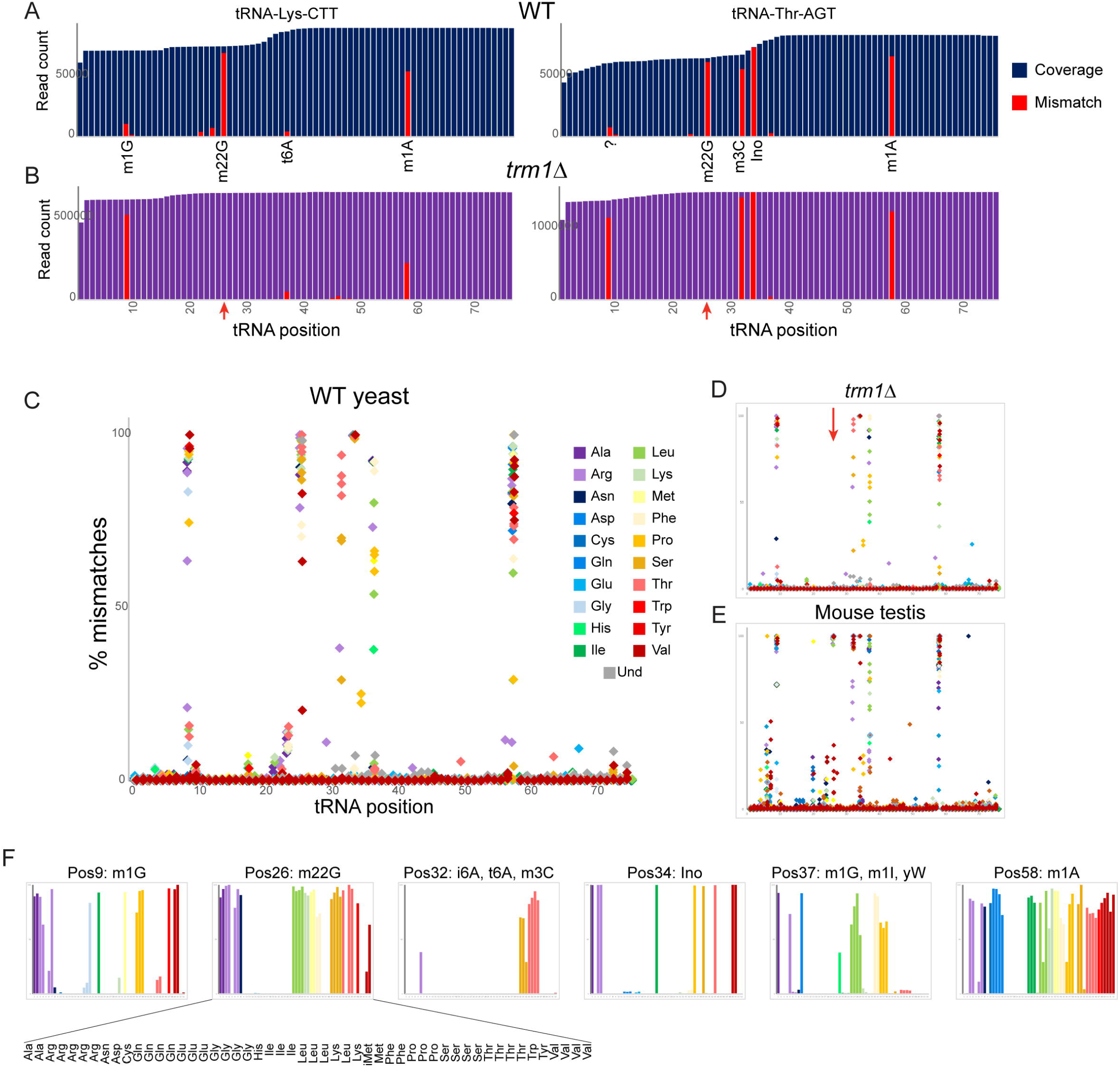
Nucleotide modifications revealed by genomic mismatches. A) Sequence coverage of the two indicated tRNAs, with reads matching the genomic tRNA sequence shown in blue, and misincorporations in red. Known nucleotide modifications are shown below each mismatch location. Question mark at position 9 for Thr-AGT indicates no annotated modification at this site in the MODOMICS database (although this site is a common site for the m^1^G modification in other tRNAs); interestingly, the adjacent nucleotide (position 10) is a known site for N2-methylated guanine in this tRNA. B) Misincorporation data from the *trm1*Δ dataset for the same two tRNAs as in panel (A). Red arrows indicate loss of mismatches at position 26 in both tRNAs. C-E) Frequency of mismatches across all tRNAs for wild-type yeast (C), *trm1*Δ yeast (D), and mouse (E) OTTR tRNA datasets. In each plot, the % misincorporation is shown for each tRNA position (x axis) for each tRNA species (indicated by colors) with over 2000 reads. Red arrow in panel (D) shows the loss of the misincorporation signature at position 26 in *trm1*Δ yeast. F) Detailed view of mismatch rates for individual tRNA species at the indicated nucleotide positions from budding yeast. Bar graphs show the same data as in panel (C), zoomed in to allow easier distinction between individual tRNA species. Above each position, known nucleotide modifications found in the tRNAs exhibiting >10% misincorporation are annotated.

To explore the effects of modified nucleotides on RT misincorporation globally across all tRNA species and all positions, we plotted tRNA “mutations” for each nucleotide position across all tRNAs in wild type and *trm1*Δ yeast, and in mouse testis (**Figure 2C-E**). Consistent with prior observations [25, 27], we observed high levels of misincorporation at tRNA positions 9, 26, and 58, corresponding to well-known locations of m^1^G, m^2^_2_G, and m^1^A, respectively. Moreover, as seen for individual tRNAs (**Figure 2B**, red arrows) we find that the mutational signature at position 26 is completely lost in the *trm1*Δ background (**Figure 2D**, red arrow), confirming the causal link between the m^2^_2_G modification and the misincorporation signature at this position. Closer examination of the specific tRNA species exhibiting mismatches at any given position in yeast (**Figure 2F**) confirmed that mismatches were only observed in the subset of tRNAs known to be modified at the position in question [33], as for example alanine tRNAs carry m^1^G at position 9 whereas histidine tRNAs do not (**Figure 2F**, leftmost panel).

Beyond the mutational signature observed at known locations of m^2^_2_G, m^1^G, and m^1^A, several other locations were associated with sequencing mismatches (**Figures 2C,F**). These included known locations for N6-threonylcarbamoyladenosine (32), N6-isopentenyladenosine (32), 3-methylcytidine (32, position 20 in mouse), inosine and 1-methylinosine (34 and 37), and wybutosine (37), as well as lower misincorporation rates at several locations currently annotated as unmodified nucleotides in the MODOMICS database [33].

Taken together, these data demonstrate the utility of OTTR for analysis of a range of nucleotide modifications.

### Analysis of tRNA cleavage in budding yeast following RNY1p nuclease overexpression

Turning to analysis of smaller (<40 nt) RNAs, we next set out to benchmark several small RNA cloning protocols in the experimentally-tractable budding yeast model system. tRNAs can be cleaved by a number of different nucleases, including RNase A, T, and L family members, in a variety of species [4, 6, 7, 34, 35]. Conveniently, budding yeast do not encode any RNase A family members, and encode a single RNase T2 family member, RNY1p. As RNY1p overexpression has previously been reported to drive high levels of tRNA cleavage [35], we generated a construct bearing *RNY1* under the control of the galactose-inducible p*GAL1-10* promoter. We confirmed by Northern blot analysis that overexpression of RNY1p lead to high levels of tRNA-Gly-GCC cleavage in our hands (**Figure 3A**), providing a convenient system for production of high levels of tRNA fragments for benchmarking small RNA cloning protocols.

**Figure 3.**
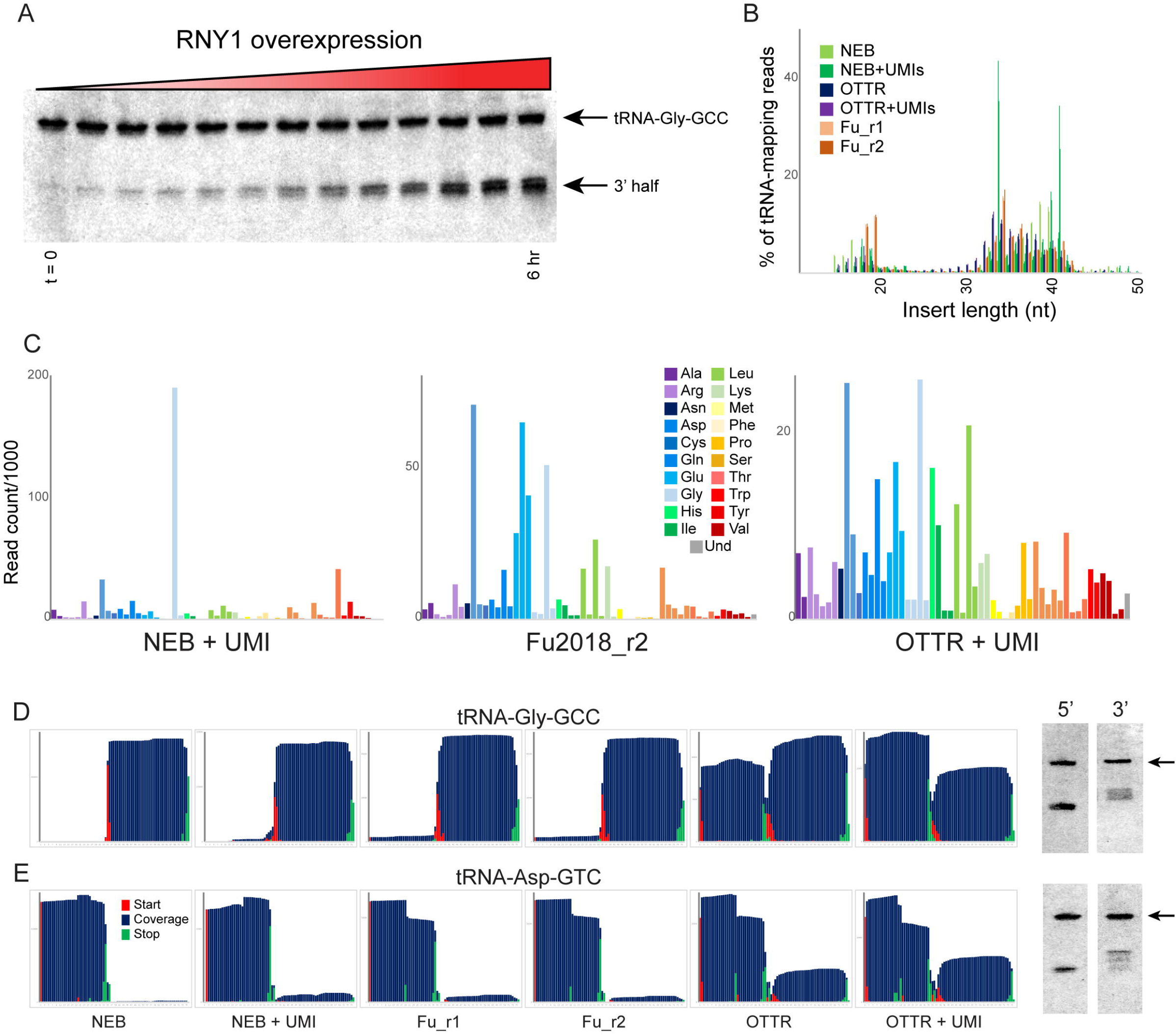
Benchmarking OTTR capture of tRNA fragments in budding yeast overexpressing RNY1p. A) Northern blots for tRNA-Gly-GCC 3’ end during a time course of RNY1p overexpression (from uninduced to 6 hour induction) in budding yeast. B) Size distributions for tRNA-mapping reads in various small RNA libraries prepared from yeast following six hours of RNY1p overexpression. See also **Supplemental Figures S3-4**. C) Overall coverage of all tRNA isoacceptors – calculated by summing all reads mapping to a given tRNA species – shown for the indicated small RNA cloning protocols. D-E) Left panels show coverage maps for tRNA-Gly-GCC (D) or tRNA-Asp-GTC (E) for the six indicated cloning protocols. Each plot shows the distribution of all 5’ (start) and 3’ (end) ends of the relevant sequencing reads, as well as the cumulative sequencing coverage across the tRNA. Right panels for each tRNA show Northern blots for the 5’ side, and the 3’ side, of the relevant tRNA (from yeast subject to six hours of RNY1p overexpression), as indicated. Black arrow highlights the full length tRNA band. For the deep sequencing dataset NEB and Fu2018 [36] protocols capture only the 3’ half of tRNA-Gly-GCC, and the 5’ half of Asp-GTC, while OTTR captures both 5’ and 3’ halves. In both cases Northern blots confirm the validity of the OTTR dataset, with both 5’ and 3’ halves present at similar abundance for both of these tRNAs.

To compare small RNA cloning protocols, we overexpressed *RNY1* for six hours in large cultures, purified total RNA, and split the RNA into aliquots for cloning. We generated an initial dataset to enable comparisons between three basic protocols: 1) a ssRNA ligase strategy developed by Fu et al [36], based on RNA ligase-dependent adaptor ligation strategies common to most small RNA-Seq protocols; 2) the NEBNext Small RNA kit; and 3) OTTR. Notably, all three protocols captured a dramatic increase in abundance of tRNA and rRNA fragments, and a shift in the size distribution of inserts, following RNY1p overexpression (**Figure 3B, Supplemental Figure S3**). All three protocols captured tRNA fragments of similar lengths, albeit with moderate differences between the three protocols – NEBNext was a particular outlier in this regard, with clear peaks of tRNA fragments that were far more prominent in these libraries than in the other two protocols.

To compare these cloning protocols in more granular detail, we calculated the representation of all yeast tRNAs in each of the various deep sequencing datasets. As shown in **Figure 3C** and **Supplemental Figure S4A**, we capture relatively few distinct tRNA fragments using the various permutations of the NEB protocol, contrasting with the far wider range of tRFs captured using the Fu2018 protocol and OTTR. Overall, we find that OTTR exhibiting the greatest diversity of tRNA fragments of all the protocols examined. We extended this analysis by binning tRNA-mapping reads according to the coverage of either the 5’ or 3’ half of each yeast tRNA. This analysis (**Supplemental Figure S4B**) again reveals remarkably few tRNA species efficiently captured by NEB Next, contrasting with the far more even coverage of tRNA fragments in our OTTR datasets (and the intermediate behavior of the Fu2018 libraries). The relatively even representation of yeast tRNA species, and 5’ and 3’ halves, is most consistent with the expectation that RNase T2 family members like RNY1p should have only modest sequence preferences beyond a preference for pyrimidines present in single-stranded RNA loop regions, and should therefore cleave most tRNAs with roughly similar efficiency. Moreover, the relatively even distribution of ratios between 5’ and 3’ halves across all tRNAs observed in the OTTR datasets is consistent with Northern blot results (see below) demonstrating roughly similar levels of 5’ and 3’ cleavage products for all four tRNAs assayed.

To enable validation by comparison to an independent measure of tRNA fragment levels, we next examined nucleotide-resolution coverage data for several tRNAs that exhibited substantial differences in abundance between these three protocols. **Figures 3D-E** show coverage plots for two exemplar tRNAs – chosen based on dramatic differences in capture the two halves of the tRNA across the three different protocols – with coverage of the tRNA shown in blue along with the locations of read 5’ (start) and 3’ (end) ends. For example, although all three protocols robustly captured the 3’ half of tRNA-Gly-GCC, OTTR uniquely captured a similar abundance of 5’ fragments of this tRNA that were absent from the other two libraries (**Figure 3D**). To validate these protocols by comparison to an independent “ground truth,” we assayed the 5’ and 3’ halves of four tRNAs by Northern blotting (**Figures 3D-E**, right panels, and **Supplemental Figure S5**). For the two tRNAs for which the deep sequencing protocols showed substantial differences in tRNA coverage, OTTR more faithfully captured the tRNA fragments observed in the Northern blots, thus validating this protocol as a more faithful representation of tRNA fragment abundance in vivo.

Taken together, our data show that OTTR captures a wider range of tRNA fragments, more even levels of 5’ and 3’ tRNA halves, and better agrees with ground truth Northern blots, all of which strongly support the utility of OTTR for analysis of tRNA-derived small RNAs.

### The small RNA payload of mature mouse spermatozoa

We finally turned to analysis of small RNAs in the mouse germline. Scores of studies over the past decade have documented abundant tRNA fragments in mammalian sperm, with the majority of published datasets documenting highly abundant 5’ tRNA fragments, with the 5’ halves of tRNA-Glu-CTC, tRNA-Val-CAC, and tRNA-Gly-GCC representing the three most abundant tRNA fragments in mouse sperm [30, 37]. However, it has been clear for years that standard deep sequencing analyses are insufficient to fully capture the sperm RNA payload. First, RNA cleavage by RNase A and T1/T2 family members is known to leave RNA 3’ ends bearing a cyclic 2’-3’ phosphate (which can spontaneously resolve to 2’ or 3’-phosphorylated ends), a modification that prevents RNA ligation during cloning. Indeed, resolving cyclic 2’-3’ phosphates via PNK treatment [38] resulted a dramatic shift in captured sperm RNAs [16], revealing a far greater abundance and diversity of rRNA cleavage products than previously appreciated, along with longer 5’ tRNA fragments that presumably reflect the primary cleavage site for reproductive tract nucleases. Second, Northern blotting studies in sperm and epididymis samples revealed the presence of 3’ tRNA halves that are not represented in deep sequencing datasets [16, 17], further emphasizing our incomplete understanding of the mammalian sperm RNA payload.

To directly compare small RNA cloning protocols on mouse sperm RNAs, we pooled cauda epididymal sperm from ten males for total RNA extraction. Total RNAs were *mir*Vana size selected prior to being split into three large aliquots and either 1) left untreated, 2) treated with PNK in the absence of ATP to drive 3’ end dephosphorylation, or 3) treated with PNK and ATP to both resolve 3’ phosphates and to phosphorylate RNA 5’ ends. Each pool was then further split into three aliquots and cloned using either Illumina TruSeq, NEBNext Small RNA, or OTTR. **Figures 4A-C** show insert size distributions, mapping rates to various RNA species, and abundance of various tRNA species, respectively.

**Figure 4.**
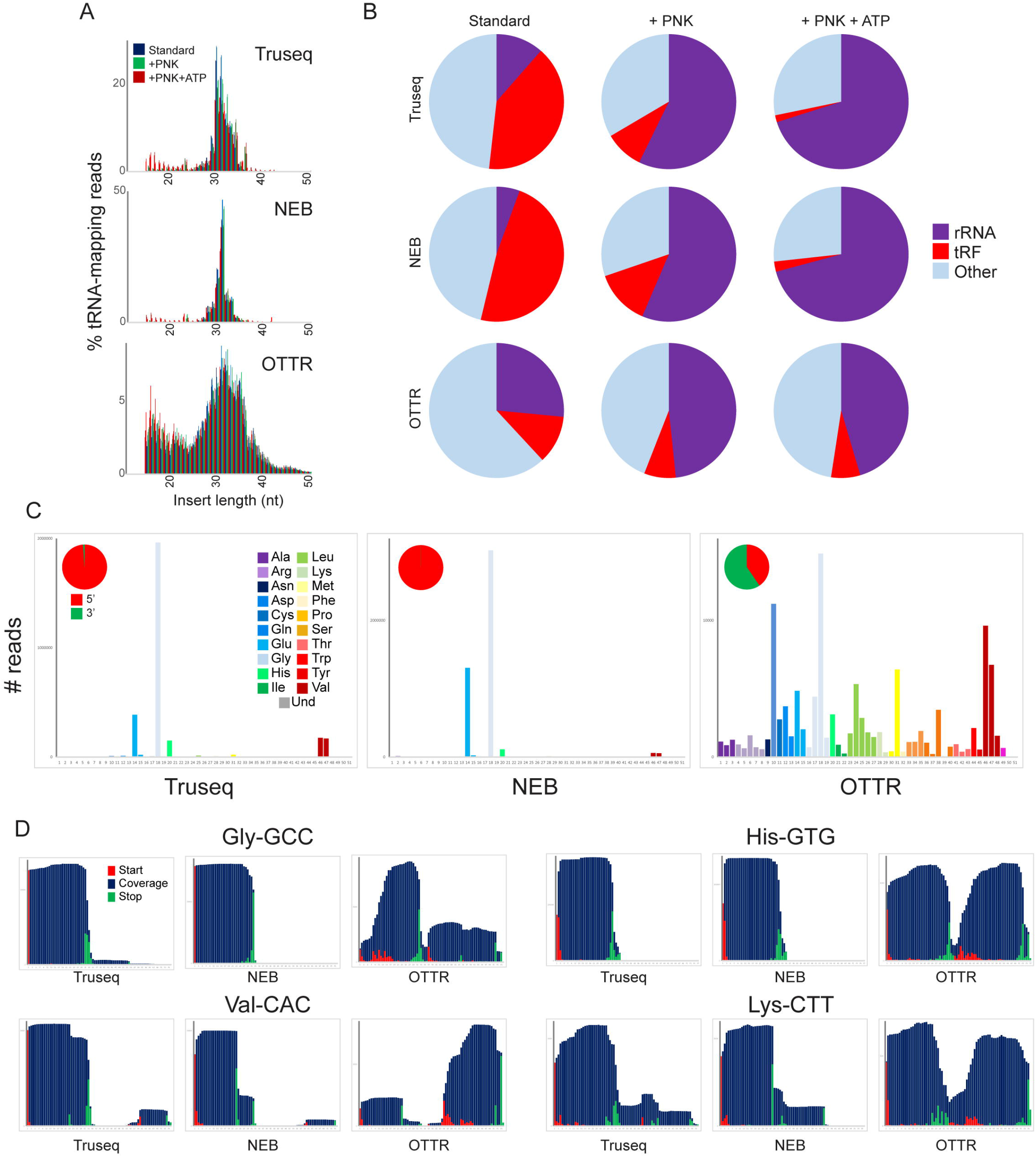
A revised view of the mouse sperm small RNA payload. A) Small RNA length distributions, as in **Figure 3B**, for tRNA-mapping reads in various mouse sperm RNA libraries generated using the indicated protocols. B) Pie charts showing overall mapping of each library to the indicated RNA classes. C) Overall coverage of all tRNA species in each dataset, as in **Figure 3C**. Here, each dataset also has a pie chart showing the percentage of tRNA-mapping reads derived from the 5’ or the 3’ half of tRNAs, as indicated. See also **Supplemental Figure S6**. D) Coverage plots for four typical tRNAs, as in **Figures 3D-E**.

Focusing first on ligation-based small RNA libraries, our untreated TruSeq and NEBNext datasets recapitulated features of mouse sperm RNAs documented in many prior TruSeq studies [30, 37], including abundant 5’ tRFs deriving primarily from tRNA-Glu-CTC, tRNA-Val-CAC, and tRNA-Gly-GCC (**Figure 4C, Supplemental Figure S6**). In contrast, we find that OTTR reveals a far greater range of tRNA fragments than either of the commercial protocols. Consistent with our findings in yeast, OTTR libraries capture a much broader representation of tRNAs than either of the ligation-based protocols, which are dominated by the handful of abundant tRFs – Glu-CTC, Gly-GCC, Gly-CCC, Val-CAC – seen in many prior studies. Moreover, while the ligation-based tRNA reads almost exclusively map to tRNA 5’ ends, OTTR libraries carry both 5’ and 3’ fragments of most tRNAs (**Figures 4C-D**). These findings are consistent with prior Northern blot studies demonstrating the presence of both 5’ and 3’ tRNA halves in mouse sperm, again supporting the broad utility of OTTR for tRNA fragment cloning.

We finally asked whether 3’ phosphate modifications impact the ability of OTTR to capture various small RNA species in mouse. Focusing first on the ligation-based cloning methods, we confirm prior reports [16, 39, 40] showing that PNK treatment (with or without ATP) resulted in a massive increase in capture of rRNA-derived fragments (**Figure 4B**), again consistent with the hypothesis that rRNA fragments in mammalian sperm are generated by a nuclease of the RNase A, T, or L families. PNK treatment also enabled improved capture of specific cleavage products in the TruSeq and NEB libraries, as assessed by increased overall levels of particular tRNA fragments (**Supplemental Figure S7**) and increased capture of specific 3’ ends (**Supplemental Figure S8**).

Compared to the ligation-based small RNA-Seq libraries, OTTR libraries were far less affected by PNK treatment, as OTTR libraries prepared from untreated sperm RNAs exhibited similarities to ligation-mediated libraries prepared from PNK-treated samples. This is illustrated in the mapping rates to different RNA species, where untreated OTTR libraries exhibited similar levels of rRNA cleavage products as the PNK-treated TruSeq or NEBNext libraries, and were only modestly affected by PNK treatment (**Figure 4B**). Similarly, many of the tRNA fragments that required PNK treatment for capture in TruSeq or NEB libraries were already abundant in untreated OTTR libraries and again unaffected by PNK treatment (**Supplemental Figure S7A**). Nonetheless, PNK treatment did lead to improved capture of a subset of tRNA fragments in OTTR libraries (**Supplemental Figure S7A**), and close examination of 3’ cleavage sites revealed capture of longer species for some 5’ tRFs (see for example **Supplemental Figure S7B, red arrows**). Taken together, our findings demonstrate that although OTTR appears to be able to capture a subset of small RNAs bearing 3’ phosphates and/or 2’-3’ cyclic phosphate moieties, PNK treatment nonetheless enhances capture of some RNA cleavage products by this protocol and thus should be included for analyses of tRNA and rRNA cleavage.

## DISCUSSION

Here, we compared the performance of a variety of small RNA cloning protocols in characterization of tRNAs and tRNA fragments. By several metrics, we find that OTTR outperforms major commercial RNA-Seq protocols, and performs comparably to the mim-tRNAseq and YAMAT-Seq protocols [25, 26] designed for tRNA cloning. OTTR efficiently captures full-length tRNAs without requiring enzymatic removal of nucleotide modifications, with fewer than 10-25% of tRNA sequences (depending on the species under study) suffering from premature termination (**Figures 1C-D**). Moreover, as seen in other tRNA cloning datasets, nucleotide misincorporation events provide a readout of a number of nucleotide modifications, including m^1^G, m^2^_2_G, m^3^C, m^1^A, i6A, t6A, inosine, and wybutosine.

Not only does OTTR accurately capture tRNAs with fidelity similar to that of protocols specifically optimized for tRNA sequencing, OTTR was previously shown to exhibit the best performance in small RNA sequencing when benchmarked using miRXplore – a synthetic miRNA reference standard of 962 small RNA oligos [28]. Here, we additionally show that OTTR more accurately captures tRNA fragments than three typical small RNA-Seq protocols, again highlighting the range of applications for this cloning protocol. Taken together, the ease of library preparation, along with the documented performance of OTTR in both microRNA and tRNA cloning applications, highlight the diversity of scenarios for which OTTR is an ideal small RNA cloning protocol.

A variety of enzymatic or chemical RNA treatments can be envisioned that would modify the range of RNAs captured by OTTR. Most notably, given the enhanced capture of full length tRNAs observed in the absence of m^2^_2_G in RNA from *trm1*Δ yeast, applications focused on intact tRNA capture would presumably benefit from AlkB-mediated tRNA demethylation – as pioneered in the ARM-Seq and DM-Seq methods [22-24] – to remove modifications that interfere with RT processivity. The drawbacks to this protocol modification include the potential for tRNA fragmentation resulting from AlkB treatment, presenting challenges to subsequent full length tRNA capture, as well as loss of the ability to assay nucleotide modifications by nucleotide misincorporation at these sites (**Figure 2**). OTTR could also be modified to characterize tRNA charging status by treating total RNA to maintain tRNA aminoacylation (isolation of total RNA under acidic conditions), deacylate all tRNAs (isolation under basic conditions), or specifically capture acylated tRNAs (3’ nucleotide removal from unacylated tRNAs via periodate oxidation and β-elimination, followed by base treatment to deacylate remaining charged tRNAs to enable their subsequent cloning). These and other modifications may prove beneficial depending on the goals of any particular study.

### Revisiting the mouse sperm RNA payload

Biologically, our primary interest in benchmarking OTTR was to further explore the still-mysterious small RNA composition of mammalian sperm populations. Over the past decade, scores of studies have largely agreed in defining the mammalian sperm small RNA payload as being dominated by 5’ tRNA fragments, with 5’ ends of tRNA-Glu-CTC, Val-CAC, Val-AAC, Gly-GCC, and Gly-CCC being most abundant [30]. rRNA fragments have also been highlighted in several studies, although in our experience rRNA fragments proved the most variable between experimentalists, raising the concern that levels of rRNA fragments might be particularly susceptible to artifacts arising during cell lysis, RNA isolation, or library preparation.

For several years, it has been clear that the consensus view of mammalian sperm RNAs has been incomplete. We previously showed that removal of 3’ phosphate or 2’-3’ cyclic phosphate modifications revealed a massive population of rRNA fragments, as well as slightly longer 5’ tRNA fragments than typically captured, consistent with RNase A or T family cleavage events being responsible for rRNA and tRNA cleavage in the germline [16]. Moreover, several groups used Northern blots to show that 3’ tRNA fragments are in fact present in sperm, despite their absence from small RNA-Seq datasets [16, 17].

Here, we build on these studies using OTTR to provide the most accurate picture of the sperm small RNA payload to date. We find that sperm carry a population of small RNAs dominated by rRNA fragments, along with both 5’ and 3’ tRNA halves arising from the majority of tRNAs. Smaller populations of microRNAs and piRNAs are also present, consistent with prior reports of the sperm RNA payload. This revised view of the sperm RNA payload raises two major biological questions.

First, our findings undermine the view of sperm RNAs based on the privileged abundance of 4-5 specific tRNA 5’ halves, where only specific tRNAs are subject to cleavage, or specific tRNA halves are stabilized and/or selected for trafficking to sperm, thus forming a special population of small RNAs for delivery to the zygote. Instead, our revised view of sperm small RNAs is more consistent with a generalized cleavage of the RNA populations of any typical cell, with rRNA and tRNA fragments being found roughly in proportion to the abundance of the intact precursor species in developing sperm or typical somatic tissues. Our data do not address the question of whether a given rRNA or tRNA fragment derives from “in situ” cleavage of rRNAs or tRNAs present at the completion of spermatogenesis – as opposed to their being generated in somatic support cells in the reproductive tract [41] – but our data motivate a reappraisal of the biogenesis of structural RNA fragments in the male germline.

Second, our data raise questions about the biochemical nature of the RNAs delivered to the zygote upon fertilization. Regulatory functions have been identified for multiple solitary 5’ or 3’ tRNA fragments in isolation [8, 10, 42-45], suggesting numerous potential regulatory roles for sperm-delivered tRFs in the early embryo. However, given our finding that for many tRNAs both 5’ and 3’ tRNA halves can be found in sperm at similar abundance, it will be important to understand the biochemical context in which these RNAs are delivered to the zygote. Are both 5’ and 3’ tRNA halves still associated with one another as part of a nicked tRNA, or are the two halves dissociated and either folded into alternative conformations [46], or bound to RNA-binding proteins?

Understanding how tRNA halves are delivered has major implications for their potential functions in the early embryo.

Together, our data significantly update our understanding of the sperm epigenome, and motivate re-appraisal of mammalian germline small RNA biogenesis, and of stress and diet effects on sperm RNA populations.

## ACKNOWLEDGEMENTS

We thank P. Zamore and I. Gainetdinov for the generous gift of primers and technical assistance with the Fu 2018 ligation-based RNA cloning protocol, and R. Flynn for critical reading of the manuscript and insightful discussions. This work was funded by NIH R01HD080224 and HD099816 (HTG, OJR), F31HD097928 (CG), and the Bakar Fellows program (KC).

## MATERIALS AND METHODS

### Ethics statement

Animal husbandry and experimentation was reviewed, approved, and monitored under the University of Massachusetts Medical School Institutional Animal Care and Use Committee (Protocol ID: A-1833-18).

### Mouse husbandry and tissue collection

All samples were obtained from male mice of the FVBN/J strain background, consuming control diet Ain-93g, euthanized at 12 weeks of age according to IACUC protocol. For testis samples, both testes were collected from a single FVBN/J male, separated from the epididymis and cleaned of adhering fat, washed with PBS and snap frozen in liquid N2 for later RNA extraction.

For cauda sperm isolation, cauda epididymis samples were collected from ten males and placed into Donners complete media and tissue was cleared of fat and connective tissue before incisions were made using a 26G needle while keeping the bulk tissue intact. Tissue was gently squeezed allowing sperm to escape into solution. After incubation at 37°C for 1 hour, sperm containing media was transferred to a fresh tube and collected by centrifugation at 5000rpm for 5 minutes followed by a 1X PBS wash. To eliminate somatic cell contamination, sperm were subjected to a 1mL 1% Triton X-100 incubation 37°C for 15 mins with 1500 rpm on Thermomixer and collected by centrifugation at 5000rpm for 5 minutes. Somatic cell lysis was followed by a 1x ddH2O wash and 30 second spin 14000 rpm to pellet sperm.

### Yeast RNA purification and size selection

The yeast strains used in this study were built on the BY4741 haploid strain background according to standard methods, generating the following strains:

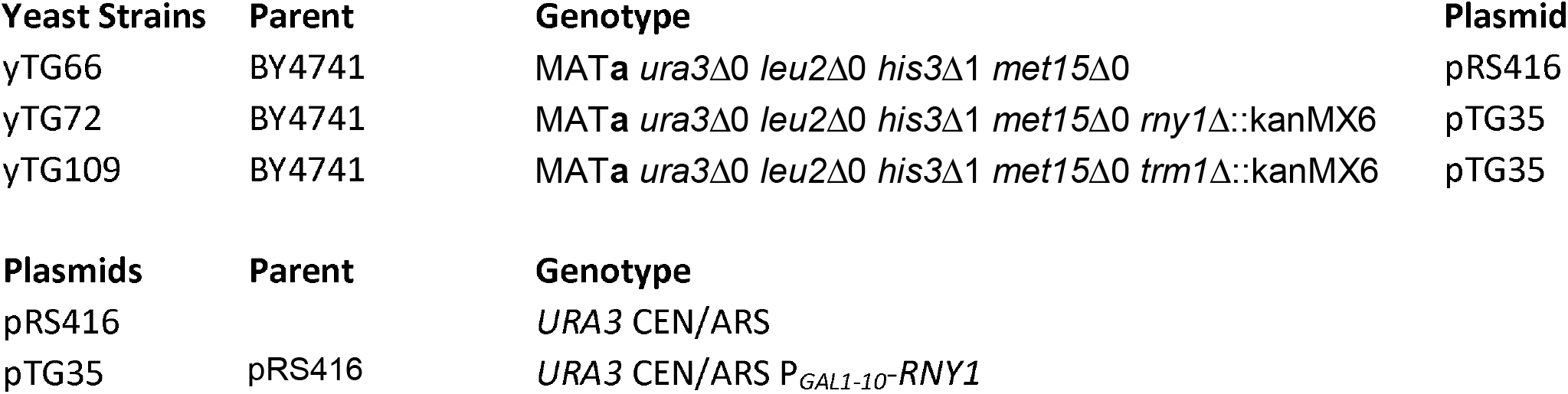

| Yeast Strains | Parent | Genotype | Plasmid |
| --- | --- | --- | --- |
| yTG66 | BY4741 | MATa *ura3Δ0 leu2Δ0 his3Δ1 met15Δ0* | pRS416 |
| yTG72 | BY4741 | MATa *ura3Δ0 leu2Δ0 his3Δ1 met15Δ0 rny1Δ::kanMX6* | pTG35 |
| yTG109 | BY4741 | MATa *ura3Δ0 leu2Δ0 his3Δ1 met15Δ0 trm1Δ::kanMX6* | pTG35 |
| Plasmids | Parent | Genotype |  |
| pRS416 |  | *URA3 CEN/ARS* |  |
| pTG35 | pRS416 | *URA3 CEN/ARS P<sub>GAL1-10</sub><sup>-</sup>RNY1* |  |

For all experiments, cells were grown at 30°C and harvested by centrifugation (2 minutes at 4000RPM in 4°C) and snap frozen in liquid nitrogen.

For full length tRNA experiments, yTG66 and yTG109 were grown overnight in selective synthetic media containing 2% dextrose, saturated cultures were diluted to OD600=0.1 and grown in selective synthetic media containing 2% dextrose until they reached OD600=0.5-07.

For tRF experiments, yTG72 were grown overnight in selective synthetic media containing 2% raffinose. Saturated cultures were diluted to OD600=0.1 in selective synthetic media containing 2% raffinose and grown until early-midlog (OD600= 0.3-0.4). Cells were then centrifuged and diluted in selective synthetic media containing 2% galactose to an OD600 so that they reach OD600=0.5-0.7 in 360 minutes (“WT” samples were resuspended in selective synthetic media containing 2% galactose, centrifuged, and snap frozen immediately).

Total RNA was prepared by resuspending cell pellets in TNE buffer (50 mM Tris-Cl pH7.4, 100 mM NaCl, 10 mM EDTA) and then vortexed with beads for a total of 2 minutes (with incubation on ice for a minute after the first minute of vortexing). Equal volume acid phenol chloroform, and SDS to a final volume of 1% was added and samples were vortexed to mix, and then incubated at 65°C for 7 minutes, followed by an additional vortex. An additional acid phenol chloroform extraction was performed followed by a chloroform extraction before RNA was precipitated, washed, and resuspended in H2O.

### Sperm RNA purification and small RNA size selection

For mouse sperm RNAs, immediately following cauda sperm purification, sperm RNAs were isolated using the *mir*Vana miRNA Isolation Kit following the enrichment procedure for small RNAs as per manual. Protocol was modified with one half volume of 100% ethanol added to the aqueous phase recovered from organic extraction (recommended volume is one third).

### Northern blots

∼3 μg of total RNA from RNY1p-expressing yeast was run on a 15% PAGE-Urea gel at 15W until dye front reached bottom of gel (∼20 min). Probes were as follows:

Arg-CCG 5’: TAACCATTGCACTAGAGGAG

Arg-CCG 3’: GCTCCTCCCGGGACTCGAAC

Asp-GTC 5’: CTGACCATTAAACTATCACG

Asp-GTC 3’: CTGACCATTAAACTATCACG

Gly-GCC 5’: TACCACTAAACCACTTGCGC

Gly-GCC 3’: GCGCAAGCCCGGAATCGAAC

Lys-CTT 5’: TACCGATTGCGCCAACAAGG

Lys-CTT 3’: GCCCTGTAGGGGGCTCGAAC

### T4 Polynucleotide Kinase (PNK) treatment

Column purified small RNAs from ten animals were pooled and split into groups: T4 PNK treatment with ATP, T4 PNK treatment without ATP, and a control no treatment group. T4 PNK treatment with ATP was incubated at 37°C for 30min in T4 PNK reaction buffer, 10mM ATP, and 50U T4 PNK (NEB) M0201S. T4 PNK treatment without ATP was incubated at 37°C for 30min in reaction buffer pH6 and 50U T4 PNK (M0201S). All sperm small RNAs samples were then cleaned and concentrated using RNA Clean & Concentrator(tm)-5 (Zymo) prior to library preparation.

### Small RNA sequencing

Small RNA sequencing was performed using one of four protocols: TruSeq Small RNA Library Preparation Kit (Illumina), NEBNext® Small RNA Library Prep Set for Illumina® (NEB), a standard in-house ligation-based cloning protocol [36], and Collins Lab OTTR Library Preparation Kit [28]. TruSeq and NEBNext library preparation was performed according to manufacturer instructions.

### Fu et al 2018 [36] ligation-based library preparation

A 3’ DNA adapter – containing an UMI sequence in 3nt blocks of random nucleotides separated by pre-defined 3nt consensus sequences, an adenylated 5’ end and a dideoxycytosine blocked 3’end – was ligated to size-selected small RNAs using T4 Rnl2tr K227Q (NEB, M0351L) for 16 hours at 25°C. The ligated product was then purified on a 10% PAGE-Urea gel, followed by gel extraction and ethanol precipitation. The purified ligated product was then ligated to a mix of equimolar 5’ RNA adaptors containing UMIs in 3nt blocks of random nucleotides separated by two distinct pre-defined 3nt consensus sequence with T4 RNA ligase (Ambion, AM2141) for 2 hours at 25°C. The final ligated product was then ethanol precipitated, and cDNA synthesis was performed with AMV reverse transcriptase (NEB, M0277L). cDNA was finally PCR amplified with standard Illumina sequencing primers with AccuPrime Pfx DNA polymerase for 12-14 cycles. Final PCR product was cleaned up with PCI extraction followed by ethanol precipitation and finally separated by 7.5% PAGE-Urea to remove adaptor dimers. The desired product was excised from the gel and eluted in 750ul elution buffer overnight at room temperature, followed by isopropanol precipitation and resuspension in 9 µl H_2_O.Final libraries were pooled and sequenced on Illumina NextSeq 500 with a 75-cycle high-output kit.

### Ordered two-template relay

OTTR was performed as described in Upton *et al* 2021 [28]. Briefly, total RNA was size selected either by mirVana (<200nt) or gel purification (60-100nt). Input RNA was labelled at the 3’ end by incubation in termal transferase buffer containing ddATP only for 90 minutes in 30°C, followed by an addition of ddGTP and another incubation at 30°C for 30 minutes. After heat inactivation of the labelling reaction (65°C for 5 minutes), unincorporated ddATP/ddGTP were hydrolyzed by incubation in 5 mM MgCl2 and 0.5 units of shrimp alkaline phosphatase (rSAP) at 37°C for 15 minutes. rSAP reaction was stopped by addition of of 5 mM EGTA and incubation at 65°C for 5 minutes. Samples were then incubated in templated cDNA synthesis buffer, adaptors, and dNTPs at 37°C for 20 minutes, followed by heat inactivation at 65°C for 5 minutes.

cDNA was size selected on an 8% PAGE-Urea gel to minimize adaptor dimers. Size selected cDNA was PCR amplified for 12-14 cycles with KAPA HiFi hot start (KAPA Biosystems, KK4602). Final PCR product was cleaned up with PCI extraction followed by ethanol precipitation and finally separated by 7.5% PAGE-Urea to remove adaptor dimers. The desired product was excised from the gel and eluted in 750ul elution buffer overnight at room temperature, followed by isopropanol precipitation and resuspension in 9µl H_2_O.Final libraries were pooled and sequenced on Illumina NextSeq 500 with a 75-cycle high-output kit.

### Extended 3’ adaptor ligation

For each of the two ligation-based protocols used for mouse sperm (Truseq, NEB Next), two replicate libraries for mouse sperm were prepared with the additional condition of an 18 hour ligation at 16°C for the ligation of the 3’ adapter in the attempt to increase ligation efficiency. For OTTR, an 18 hr incubation was added for half of libraries after terminal labeling, during the cDNA synthesis step. However, these interventions had minimal effect on sperm small RNA profiles and both replicates are interchangeable.

### Data availability

Deep sequencing data will be submitted to GEO during review and revision.

### Data analysis

To analyze small RNA sequencing data, we first removed adapters using cutadapt (version 2.9) and PCR duplicate were removed with seqkit (version 0.14.0). The trimmed and deduplicated reads were then analyzed using both an in-house pipeline and the unpublished tool tRAX (version 1.0.0; http://trna.ucsc.edu/tRAX/; [31]) and the results were compared. Firstly, we used Bowtie (version 1.1.0) to map the reads to the annotated rRNA, snoRNA, snRNA, and tRNA sequences in the corresponding species (yeast, mouse, and human) in descending priority, and then the unmappable reads to the respective genomes. The Bowtie parameters used for rRNA, snoRNA, snRNA and genome alignment were “-v 0 -k 1”; while the Bowtie parameters used for tRNA alignment were “-y -k 100 --best --strata” considering tRNA nucleotide modifications. The abundance of each type of small RNAs was normalized by the total sequencing depth, i.e., the total number of small RNA and genome mapping reads in a sequencing library. Secondly, we used tRAX to process the trimmed and deduplicated reads with default parameters; the results largely agreed with what was obtained with our in-house pipeline.

## SUPPLEMENTAL MATERIALS

## SUPPLEMENTAL TABLE LEGEND

**Supplemental Table S1. Sequencing datasets**

Deep sequencing datasets generated for this study.

## SUPPLEMENTAL FIGURE LEGENDS

**Supplemental Figure S1.**
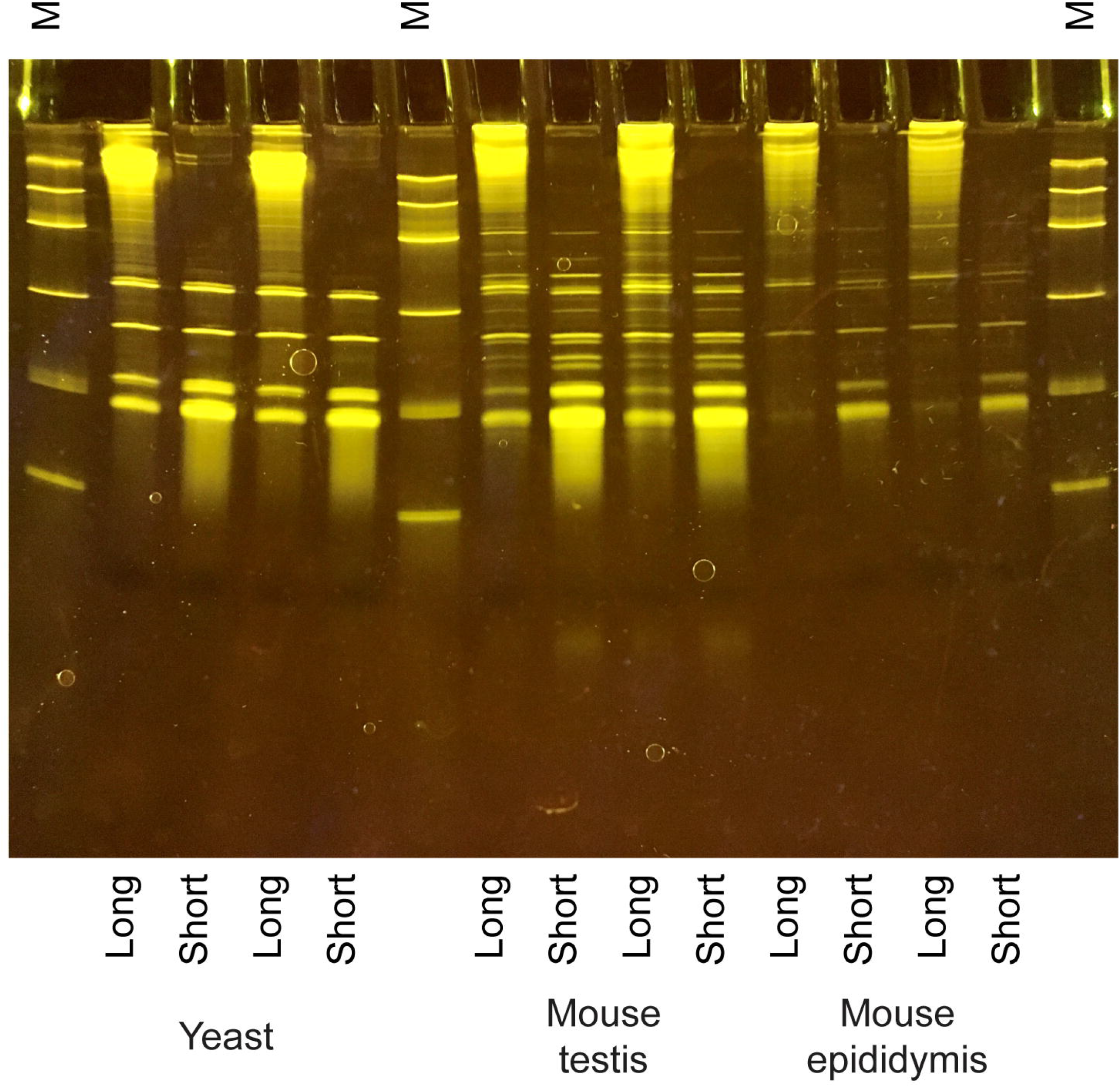
Size selection for OTTR libraries. Gel shows *mir*Vana column-enriched short and long RNA fractions, isolated from yeast and various mouse tissues, as indicated.

**Supplemental Figure S2.**
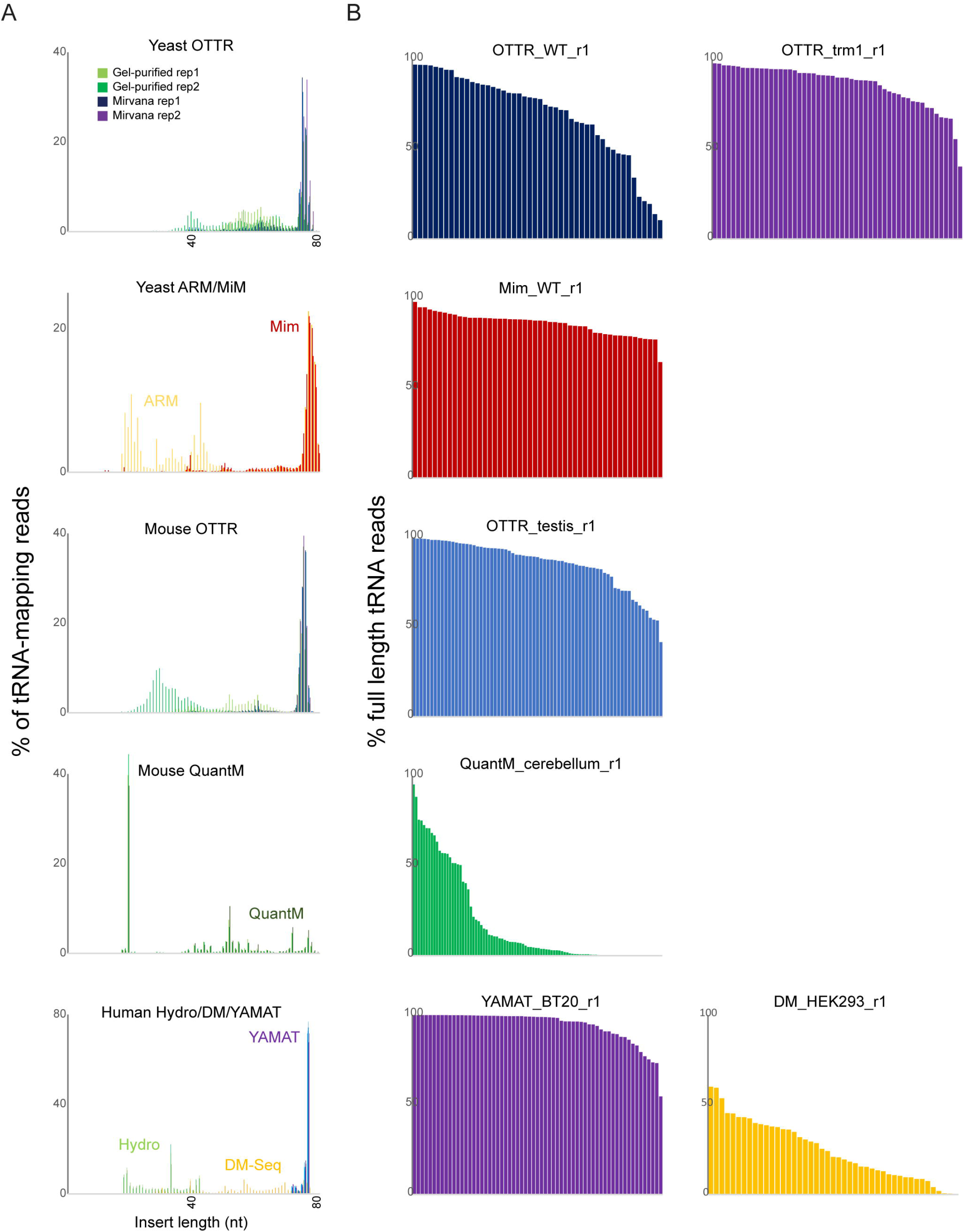
Full length tRNA capture across published protocols. A) Length distributions for tRNA-mapping reads from the indicated datasets, as in **Figure 1A**. B) For the indicated protocols, tRNAs are ordered according to % of full-length inserts, as in **Figures 1B, C**, and **E**. OTTR captures intact tRNAs with efficiency comparable to Mim-tRNAseq and YAMAT-Seq, with remaining protocols all exhibiting little to no capture of intact tRNAs.

**Supplemental Figure S3.**
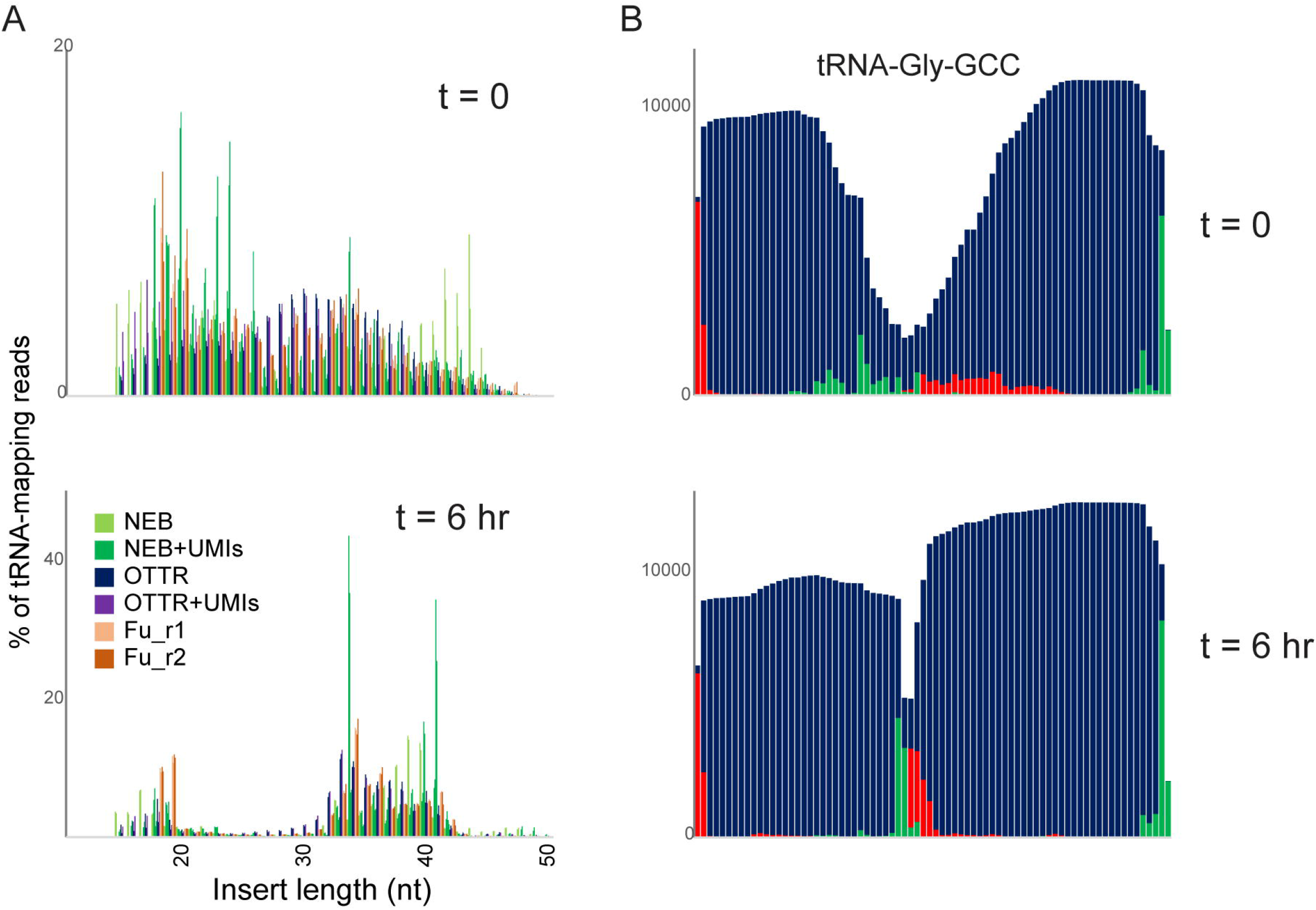
RNY1p expression drives tRNA cleavage in yeast. A) Size distribution of tRNA-mapping reads in the indicated libraries prepared from yeast carrying the p*Gal*:*RNY1* plasmid but grown under noninducing raffinose conditions (top panel), or grown for six hours in galactose to induce RNY1p (bottom panel). The variation in read start and end locations prior to RNY1p induction likely reflect nonspecific tRNA degradation during RNA handling, in contrast to the induction of precise cleavage at the anticodon following RNY1p induction. In addition, spike-ins of *S. pombe* for normalization confirm a ∼10-fold increase (not shown) in tRNA fragments following RNY1p induction in *S. cerevisiae*. B) Coverage of tRNA-Gly-GCC in OTTR libraries before (top) and after (bottom) RNY1p overexpression. As in panel (A), specific cleavage at the anticodon is readily distinguished from the more nonspecific tRNA degradation seen in uninduced conditions.

**Supplemental Figure S4.**
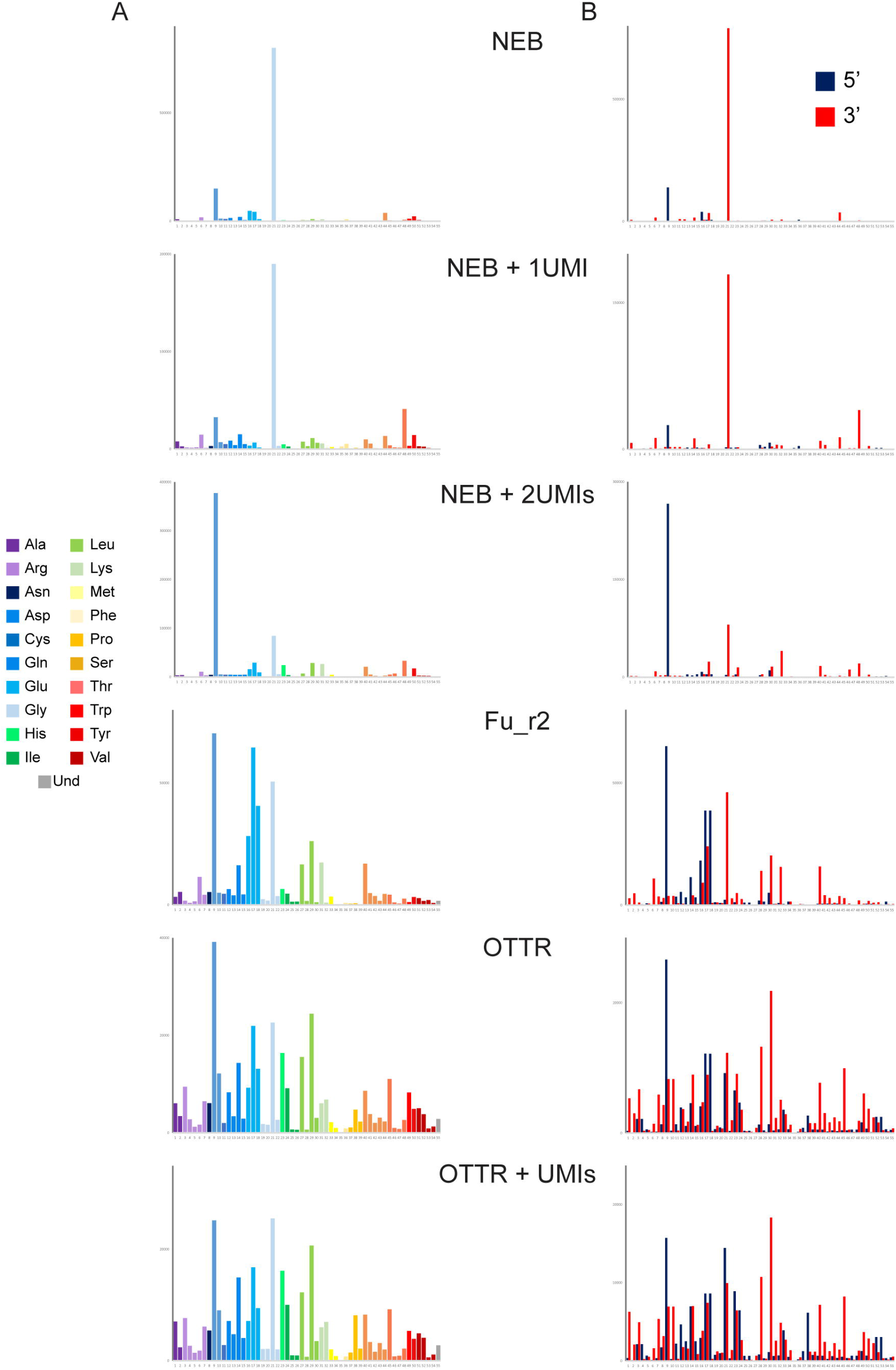
Poor capture of tRNA fragments by ligation-based methods. A) Coverage of all yeast tRNAs in RNY1p-expressing yeast. As in **Figure 3C**, for all libraries sequenced. B) 5’ and 3’ coverage. As in panel (A), but here reads were separately mapped to tRNA 5’ or 3’ halves and the two halves are plotted separately, as indicated.

**Supplemental Figure S5.**
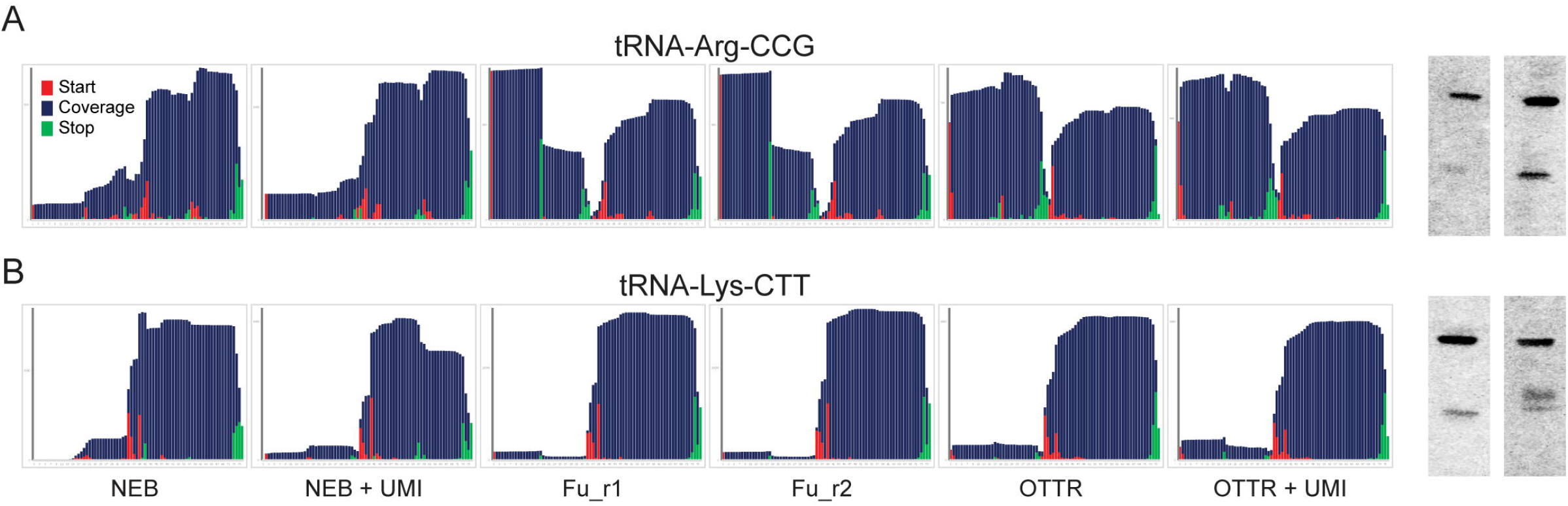
Northern blotting for tRNA halves in RNY1p-expressing yeast. As in **Figures 3D-E**, for two tRNAs which exhibited similar behavior across the six deep sequencing datasets.

**Supplemental Figure S6.**
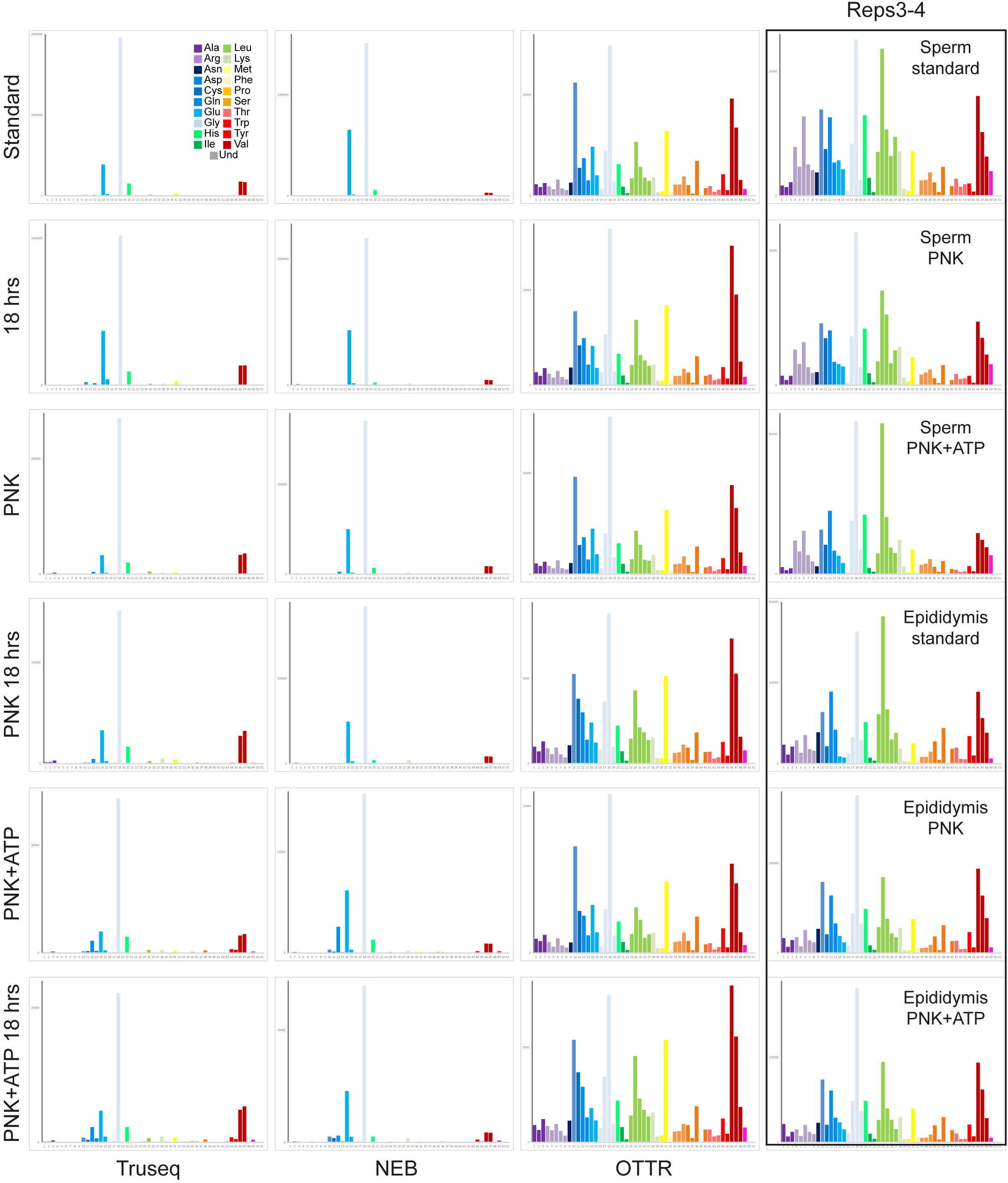
Global tRNA coverage in mouse sperm small RNAs. Left panels show coverage of all mouse tRNAs species in the indicated libraries, plotted as in **Figure 4C**. Note that for each library type we performed two biological replicate experiments; as these were essentially indistinguishable, only one replicate is shown here. The right column includes data from a second batch of OTTR datasets, in which we generated two biological replicates for mouse sperm and for mouse cauda epididymis. Again, only one of the two replicates is shown here as both replicates were nearly identical at this level of granularity.

**Supplemental Figure S7.**
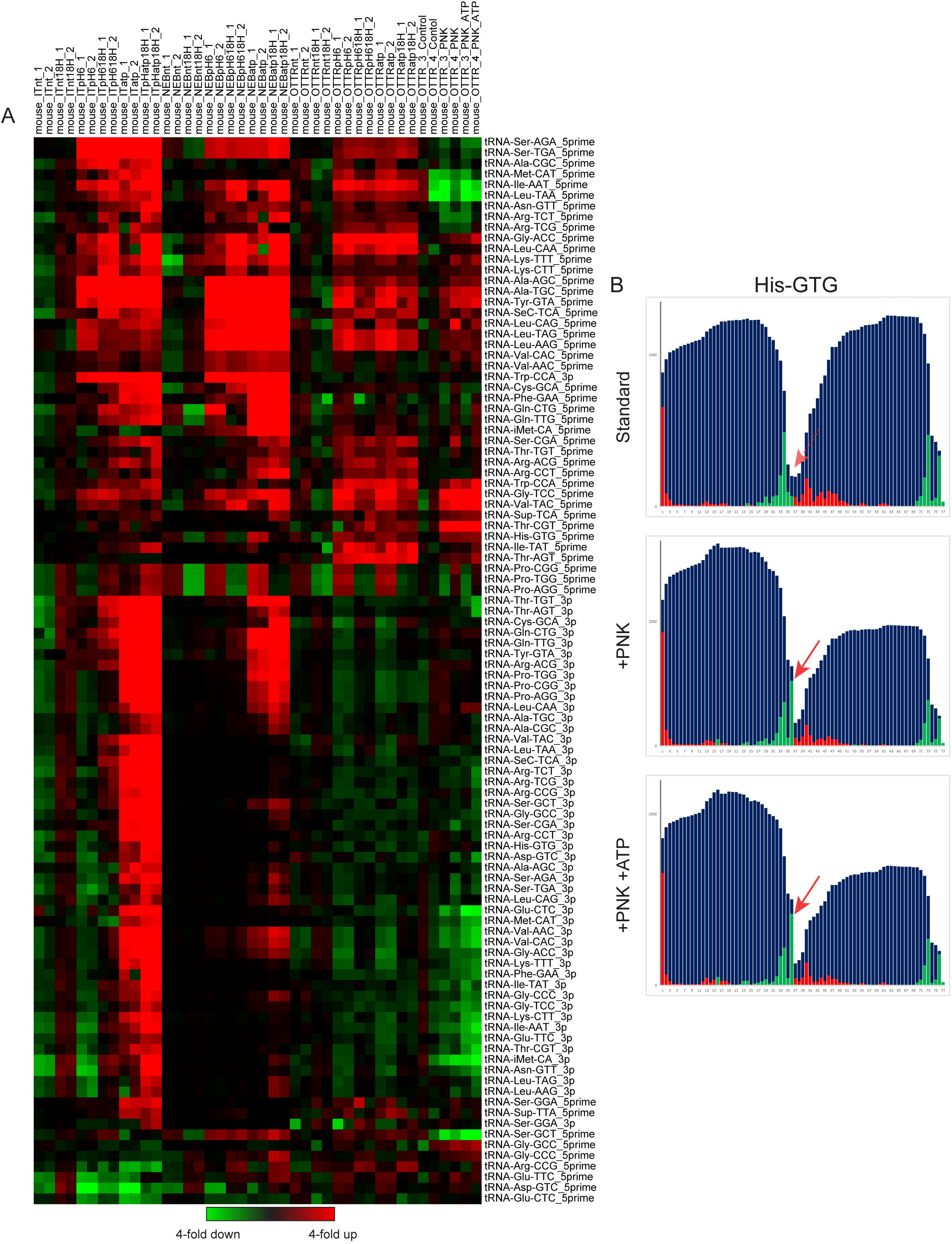
Effects of PNK treatment on tRF levels. A) For each set of libraries – TruSeq, NEB, OTTR reps1-2, and OTTR reps3-4 – relative abundance of 5’ and 3’ tRFs for each library was normalized relative to the median value across the four untreated libraries. This visualization reveals increased abundance of a wide range of tRFs resulting from PNK treatment. 3’ fragments were generally increased in abundance in TruSeq and NEB libraries following PNK+ATP treatment and were unaffected by PNK treatment of OTTR libraries, although even after PNK+ATP treatment these tRFs were still vanishingly scarce in TruSeq and NEB libraries. Many 5’ tRFs were enriched following PNK treatment in all three library conditions (eg Leu-CAA), while a smaller subset of 5’ tRFs were PNK-enriched in TruSeq and NEB datasets but were not affected by PNK treatment in the OTTR libraries (eg Cys-GCA). We initially speculated that PNK-affected and PNK-independent 3’ tRNA ends in OTTR libraries would reflect the natural 3’ ends of the tRNA fragments – the OTTR protocol begins with a nontemplated addition of A or G to small RNA 3’ ends to allow a single base pair of overlap with the C/T-ending 3’ adaptor. In this scenario, untreated RNAs with blocked 3’ ends would not be available for this single nucleotide extension, preventing cloning of tRNA fragments ending in C or T, but leaving tRNA fragments already ending with an A or G available for hybridization to the adaptor. However, close examination of PNK-dependent and -independent 3’ ends did not strongly support this hypothesis, leaving the reason for PNK effects on different tRFs unexplained at present.B.Example of PNK-dependent 3’ cleavage site capture in OTTR libraries. All three panels show read start, stop, and coverage for tRNA-His-GTG. Red arrow shows the 3’ end of a longer 5’ tRF seen only following PNK treatment.

**Supplemental Figure S8.**
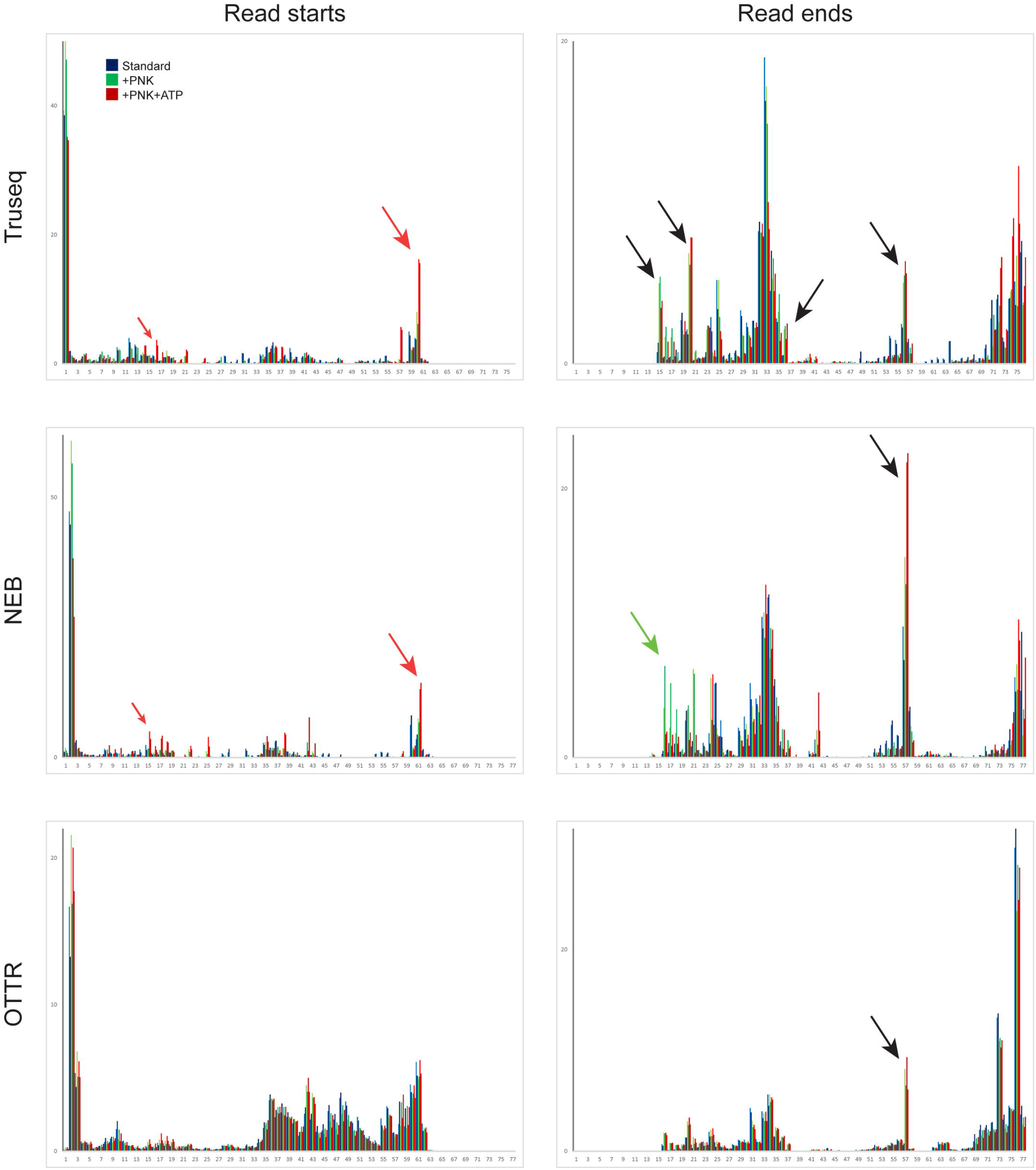
Effects of PNK treatment on tRF ends. For each tRNA in the mouse genome with over 100 reads of total coverage, the percentage of read starts (left column) or read ends (right column) was calculated for each library type. Read starts and stops percentages were then averaged across all tRNAs, and are plotted for untreated, PNK-treated, and PNK+ATP-treated libraries. Red arrows highlight PNK-specific peaks, green arrows highlight PNK+ATP-specific peaks, and black arrows highlight peaks enriched in both PNK and PNK+ATP treatments compared to untreated RNA libraries.

